# Ordered Hexagonal Patterns via Notch-Delta Signaling

**DOI:** 10.1101/550657

**Authors:** Eial Teomy, David A. Kessler, Herbert Levine

## Abstract

Many developmental processes in biology utilize Notch-Delta signaling to construct an ordered pattern of cellular differentiation. This signaling modality is based on nearest-neighbor contact, as opposed to the more familiar mechanism driven by the release of diffusible ligands. Here, we show that this “juxtracrine” property allows for an exact treatment of the pattern formation problem via a system of nine coupled ordinary differential equations. Furthermore, we show that the possible patterns that are realized can be analyzed by considering a co-dimension 2 pitchfork bifurcation of this system. This analysis explains the observed prevalence of hexagonal patterns with high Delta at their center, as opposed to those with central high Notch levels. Also, our theory suggests a simple strategy for producing defect-free patterns.

---

Biological cells can exist in a number of distinct phenotypes, even with a fixed genome. These phenotypes arise via multi-stability of the underlying dynamical network controlling cell behavior and allow cells to take on differentiated roles in overall organism function. It is clear that developmental processes must ensure that these phenotypes arise in the right place and at the right time, i.e., ensure the emergence of functional phenotypic patterns.

A well-studied case of such a system is that of Notch-Delta signaling [1]. Various cells contain Notch trans-membrane receptors [2] that couple to Notch ligands such as Delta or Jagged on both the same cell (cis-coupling) and neighboring cells (trans-coupling). Because of the specific manner by which Notch and Delta mutually inhibit each other (see below), their interaction typically leads to an alternating “salt and pepper” structure. This type of patterning is seen in systems ranging from eyes [3] and ears [4] to intestines [5] and livers [6]. As a general rule, the high Delta cells are the most specialized ones (for example, the photoreceptors [7]) and are surrounded by less differentiated high Notch supporting cells in the final structure. Parenthetically, changes in the transcriptional regulation utilizing the Delta-alternative Jagged ligand may be crucial for the role of Notch in cancer metastasis [8, 9], but here we focus solely on Delta and its interplay with Notch.

We study two aspects of the Notch-Delta system on 2d hexagonal arrays of cells. As will be seen below, this geometry allows for an exact re-writing of the (ordered) pattern-forming problem as a nine-dimensional dynamical system; analysis of this system indicates that one can understand central features of this system by expanding about a co-dimension two pitchfork bifurcation. The second aspect to be considered concerns mechanisms for ordered patterns to emerge from generic initial conditions. Here we identify a possible role for an initiating wave, similar to what has been seen in at least some biological realizations [10].

The Notch-Delta interaction is an example of juxtacrine (i.e., contact-dependent) signaling. As sketched in Fig 1, Notch ligands such as Delta bind receptors and, when this occurs between neighboring cells, leads to the cleavage of the receptor and release of its intracellular domain (NICD). This molecule translocates to the nucleus where it transcriptionally up-regulates Notch and down-regulates Delta. The ligand-receptor interaction between molecules on the same cell leads to mutual annihilation with no NICD release [11]. The combination of this cis-annihilation and the transcriptional repression is responsible for the observed lateral repression [12]. We will use a baseline model [13] of this process involving three concentrations, N (receptor), D (ligand) and I (NICD),

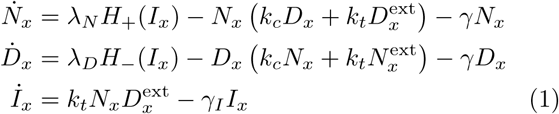

Here positions *x* refer to locations on a unit hexagonal lattice (see Fig 2) and the superscript “ext” refers the average over the six nearest neighbor sites of *x*. The production terms *H*_±_ corresponding to the aforementioned transcriptional regulation are taken to be Hill functions,

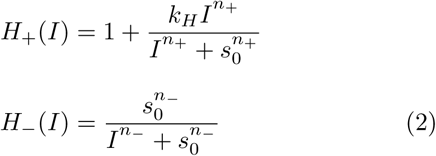

such that *H*_±_(0) = 1 and *H*_+_ is an increasing function that saturates at 1+*k*_*H*_, while *H*_-_ is a decreasing function that decays to 0 with increasing *I*. We define a typical set of parameters taken from the literature: *γ* = 0.1. *γ*_*I*_ = 0.5, *n*_+_ = *n*_-_ = 2, *k*_*c*_ = 0.1, *k*_*t*_ = 0.04, *k*_*H*_ = 1, *s*_0_ = 1, and focus on the role of *λ*_*N*_ and *λ*_*D*_.

**FIG. 1.**
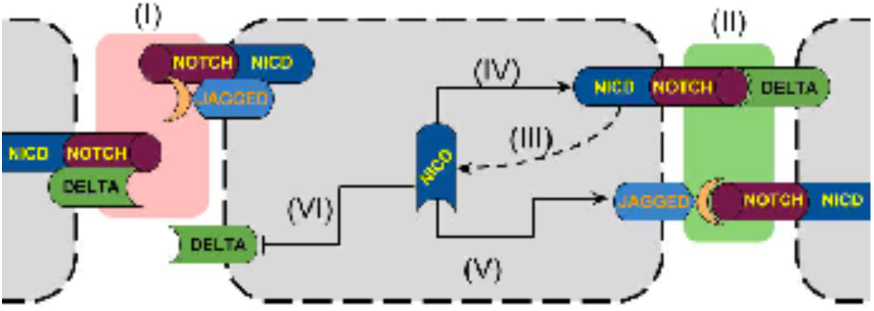
Schematic showing Delta, Jagged and Notch-NICD complexes on the cell membrane (I). Binding of Notch on one cell to Delta on the other (II) leads to the freeing of the NICD, (III), which in turns leads to the enhancement of Notch (IV) and Jagged (V) (which is irrelevant for our current concerns) and the suppression of Delta (VI).

**FIG. 2.**
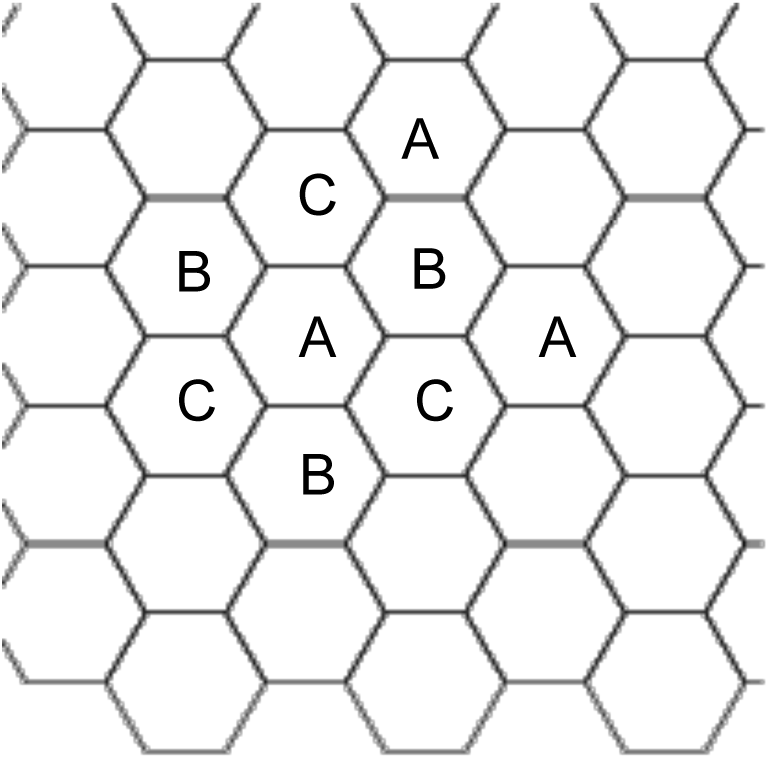
Hexagonal lattice showing the three hexagonal sub-lattices, where each A cell is surrounded by 3 B and 3 C cells, each B by 3 A and 3 C, and each C by 3 A and 3 B.

**FIG. 3.**
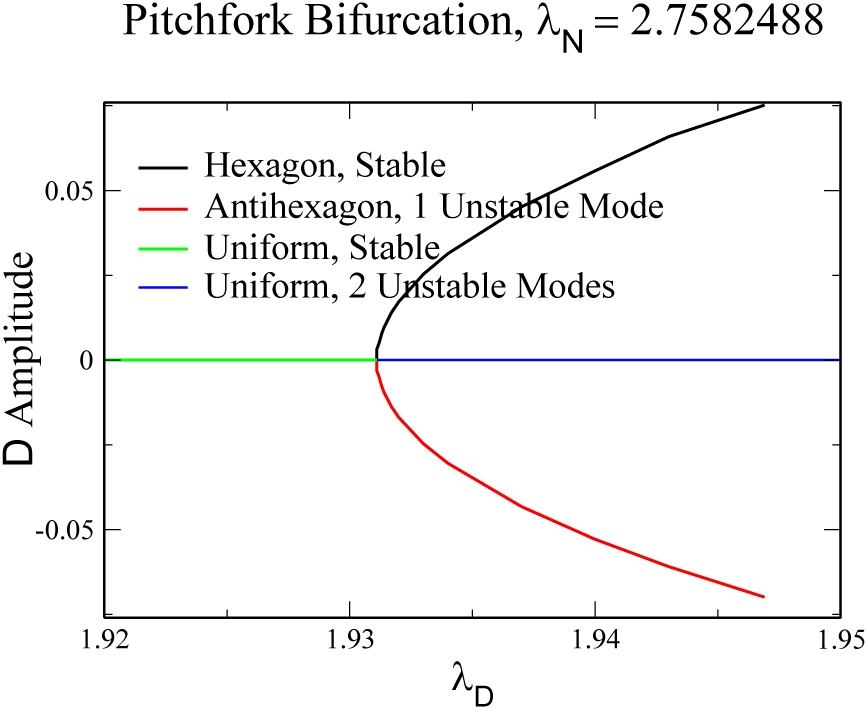
Bifurcation diagram for the critical 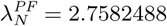 for which there is a pitchfork bifurcation.

**FIG. 4.**
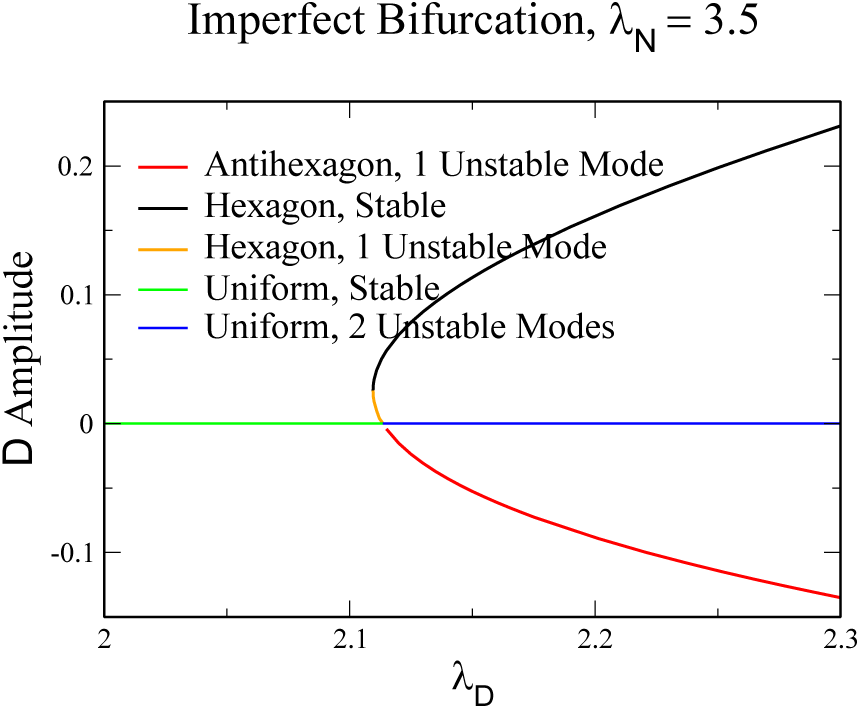
Imperfect bifurcation for 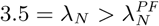.

**FIG. 5.**
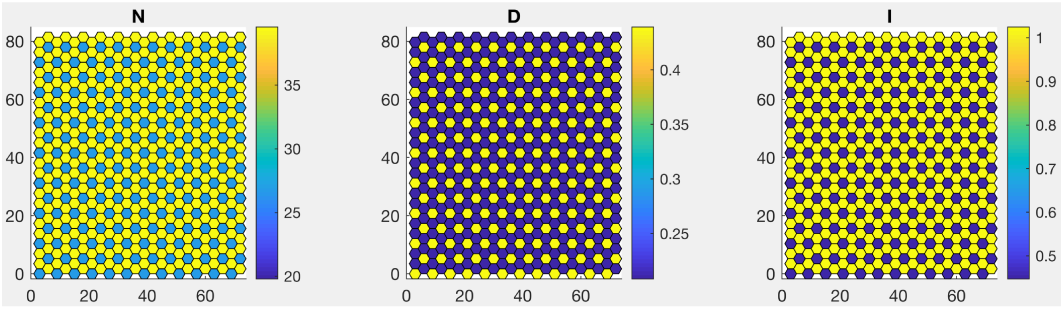
A stable hexagon solution for *λ*_*N*_ = 3.5, *λ*_*D*_ = 2.3.

**FIG. 6.**
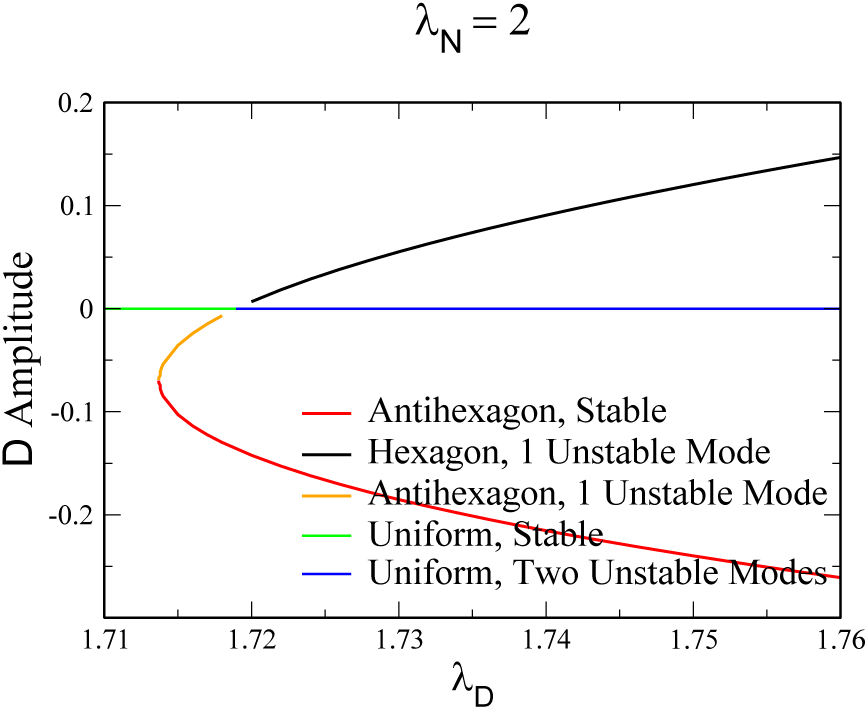
Bifurcation diagram for 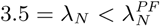.

We first consider ordered patterns; i.e., patterns that are invariant under translational invariance with vectors 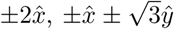. From Fig. 2, it is clear that the fields everywhere are completely determined by their values on three sub-lattices that we have labeled A, B and C. This means that the entire problem is reduced to nine coupled ODE’s. This is very different than what occurs for more traditional pattern formation problems, [14] for example for convection rolls [15], where the reduction to a set of ODE’s is valid only as an approximation near the bifurcation point. It is easy to show that at fixed *λ*_*N*_, the uniform solution with the fields taking on the same values on all three sublattices becomes unstable for 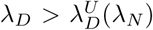 via a transcritical bifurcation, i.e. an intersection with a different nonuniform solution. On this branch, the respective values of the fields on two sublattices (say B and C) values are identical, differing from the values on the remaining (in this case, A) sublattice. This hexagonally structured solution has a 6-fold hexagonal symmetry about any site on the different (here, A) sublattice. The bifurcation is transcritical because of the lack of any symmetry between positive and negative deviations of the fields from their uniform values.

However, direct numerical solution of the ODE system show that there is a critical value of 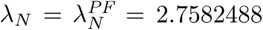 for which, due to an accidental symmetry, the hexagonally structured solution, rather than existing on either side of 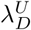, instead becomes a pair of solutions existing only for *λ*_*D*_ greater than 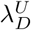, via a forward pitch-fork bifurcation. Specifically, for 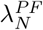, as for general *λ*_*N*_, only the uniform solution exists for 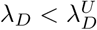 and it is stable. At 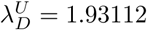, two additional solutions are born, one a “hexagon” (by definition, a solution where high *D* is surrounded by high *N*) and one an “anti-hexagon” (high *N* surrounded by high *D*). As opposed to general supercritical pitchforks, the stability of these solutions is unusual. Specifically, it can directly be shown in the 9-dimensional reduced system that the uniform state has 2 (degenerate) unstable modes above the critical *λ*_*D*_. As we will discuss below, the emerging hexagon branch is stable, whereas the anti-hexagon has one unstable mode. The instability is to a mixed mode (defined as a mode with all three sublattices having different values) which converts the anti-hexagon to a shifted hexagon.

At all other values of *λ*_*N*_, the pitchfork breaks up into a transcritical bifurcation and a saddle-node. Again by direct numerical solution, we find that for 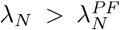, the uniform solution undergoes a transcritical bifurcation with a unstable (to a mixed-mode perturbation) anti-hexagon on the high *λ*_*D*_ side and an unstable (pure-mode) hexagon on the low *λ*_*D*_ side. The unstable hexagon then undergoes a saddle-node bifurcation, rendering the hexagon stable; this stable branch then continues on as *λ*_*D*_ increases. For *λ*_*D*_ smaller than the saddle-node value, no (anti)hexagon exists. Hence, there exists a range of parameters for which a stable hexagon coexists with the stable uniform solution, a range which widens as *λ*_*N*_ increases; we will return to this point below. For example, for *λ*_N_ = 3.5, the transcritical bifurcation in which the uniform state goes unstable is at *λ*_*D*_ = 2.11356, whereas the saddle node bifurcation is at *λ*_*D*_ = 2.1097. An example of a stable hexagon solution for *λ*_*N*_ = 3.5, *λ*_*D*_ = 2.3 is shown in Fig..

A similar thing happens for 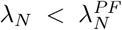, where now the stable hexagon lies to the right of the transcritical point and the unstable anti-hexagon lies to the left, and it is the one that undergoes a saddle-node bifurcation. The anti-hexagon is born with 1 unstable mode at the transcritical point and turns stable at the saddle-node bifurcation. However, unlike what happened in the previous case, the stable anti-hexagon branch loses stability to a mixed-mode perturbation; this instability leads to a pitchfork bifurcation, which is a result of the *B/C* symmetry breaking. The hexagon, on the other hand, is born with one unstable mode and subsequently becomes stable, also as a result of a mixed-mode pitchfork bifurcation. The mixed-mode solution branch arising from the hexagon bifurcation is the same solution which arises from the anti-hexagon bifurcation. For example, at *λ*_*N*_ = 2, the hexagon becomes stable at *λ*_*D*_ = 2.0477 and the anti-hexagon becomes unstable at *λ*_*D*_ = 2.056. Thus, there is a very small coexistence region between the hexagon and anti-hexagon solutions. Again, the only solution that survives stably to higher values of *λ*_*D*_ is the hexagon. This is in accord with the general rule given above that the high Delta cells are surrounded by high Notch cells sufficiently far from 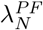 and its associated 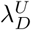.

We can use weakly non-linear bifurcation theory to make sense of these numerical findings. First, we consider parameter values away from the PF. As there are two zero modes of the uniform state at the bifurcation, there are two undetermined coefficients of the first order expansion in 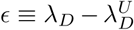, which we take to be 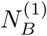 and 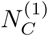, the first order shifts in *N*_*B*_ and *N*_*C*_ from the uniform solution for the shifted *λ*_*D*_. These will be fixed by higher-order terms, all of whose coefficients can be determined numerically. Doing the analysis for *λ*_*N*_ = 2.5, for example, where 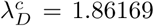, we find to second order the steady-state equations

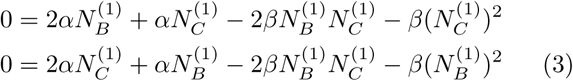

with *α* = 0.0720107, *β* = 0.000531444 and the shift in 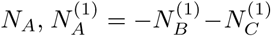. This system has four solutions

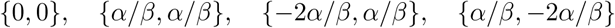

The first of these corresponds, of course, to the uniform solution, the second to the hexagon centered at *A*, the third to the hexagon centered at *B* and the last to the hexagon centered at *C*. The last three solutions are indeed hexagons for *ϵ* > 0 and antihexagons for *ϵ* < 0, since the shift in *N* at the central point is 2*α/β* < 0, i.e. a decrease in *N*, corresponding to an increase in *D* at the center, which defines the hexagon. At 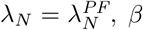 vanishes, and we have to go to third order, and for 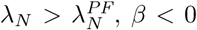 and we have hexagons for *ϵ* < 0 and antihexagons for *ϵ* > 0. From here, we also can retrieve the stability of the solutions. The stability matrix is

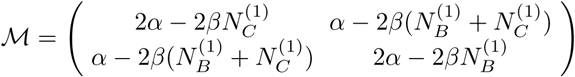

For the uniform and *A*-centered hexagon, where 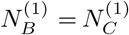, the growth rates are then

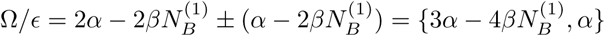

Thus, the uniform solution is twice unstable for *ϵ* > 0, whereas the hexagon has one unstable mode, as we found in the numerics. To this order, the saddle-node and the branch emerging from it are not accessible.

To see the complete structure, we consider 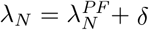. Here we have an imperfect bifurcation whose width in *λ*_*D*_ around the critical point is of order *δ*^2^, so that 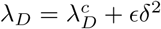. The critical point is at

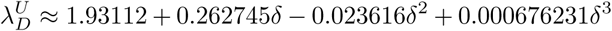

so that 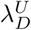 increases with *λ*_*N*_, as we have already seen. Expanding to third order in *δ*, we get the full amplitude equation

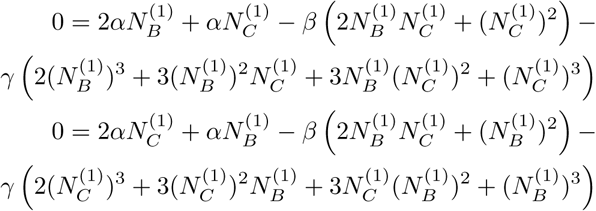

with *α* = 0.064360*ϵ, β* = 0.00166348 and *γ* = 0.00044397. To find the solutions of this system, we write 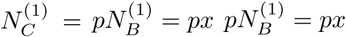. Then, the system reads

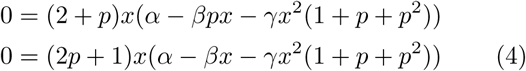

Thus, one of the factors must vanish in each of the equations. The first possible solution is clearly *x* = 0. The second solution is obtained when the last factors of both equations vanish. In this case, subtracting the two factors gives *p* = 1, in which case both equations are identical, and

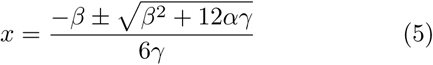

These two solutions correspond to the hexagon and antihexagon centered at *A*, emerging from the saddle-node at *α* = −(12*γ*)^-1^. Another solution is given by *p* = −2, so that the first factor of the first equation vanishes, and the setting the last factor of the second equation to zero yields again

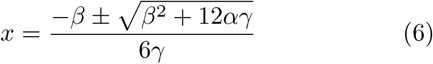

These are the hexagon and antihexagon centered at *C*. The last possibility is to choose *p* = −1*/*2 and set the last factor of the first equation to zero, so that once again

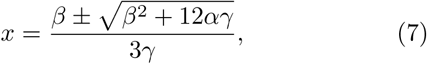

corresponding to the hexagon and antihexagon centered at *B*. We can now check the stabilities in an analogous manner and reproduce all aspects of the figures given above aside from the aforementioned mixed-mode instability. This latter structure is only visible perturbatively extremely close to 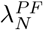, and requires a yet higher-order calculation.

The existence of stable ordered hexagon patterns leaves open the question of how these patterns can be generated in the inevitably noisy biological system with plausible initial conditions [16, 17]. In particular, it is easy to check numerically that, starting with no pattern for a set of parameters for which the uniform state is linearly unstable, the presence of noise, either in the initial data or in the time evolution, will lead to disordered states with many domain boundaries between hexagon patterns centered on different sublattices. One way out is based on the fact we have shown above that there could exist a parameter range for which there is a subcritical bifurcation to stable hexagons in which case a local perturbation which nucleates the pattern can spread in an ordered manner; this is, of course, a standard scenario in many non-living systems [14]. Intuitively, we believe that most biological systems exhibit insufficient parameter control and too high a level of stochasticity for this to be a robust strategy. There of course could be more exotic biological mechanisms that for example would provide downstream checks that prevent neighboring cells from both developing the same phenotype even if there is some initial defect in the Notch-Delta structure [18].

A more physics-based possibility is that the system is not all at once put into the unstable state. Rather, the system is initially in a regime of parameter space for which the uniform state is stable. Then some external mechanism induces a propagating wave, behind which the parameters are in the unstable region. To exhibit this possibility, we assume that only *λ*_*D*_ is affected by this wave, and *λ*_*D*_ = 2 ahead of the wave and *λ*_*D*_ = 3.5 behind the wave. We do not concern ourselves here with the origins or dynamics of this initiation wave, and rather choose a standard *tanh* waveform, and vary the wave speed *v*. In this regard, our suggestion differs from that of Ref. [[19, 20]], who start with a bistable system with two uniform states - in our proposal, the bistable dynamics is not intrinsically related to the Notch-Delta dynamics. In Fig. 7, we show a pair of simulations of our model augmented by quenched noise. At large *v*, the parameter shift is essentially instantaneous over a large spatial region and the noise nucleates incommensurate patterns in different parts of the lattice, leading to obvious defects. If the parameter wave is slowed down, the leading edge of the pattern has sufficient time to align itself with the preceding row before having to itself act as a template for the next row. In some sense, of course, all we have done is transfer the problem to one of creating a bistable system responsible for the parameter dependence. But, we assert that this is relatively easy to accomplish and that decoupling the patterning aspect from the bistable aspect (i.e., the Notch system is not bistable at the physiological parameters) is a robust approach to the elimination of defects.

**FIG. 7.**
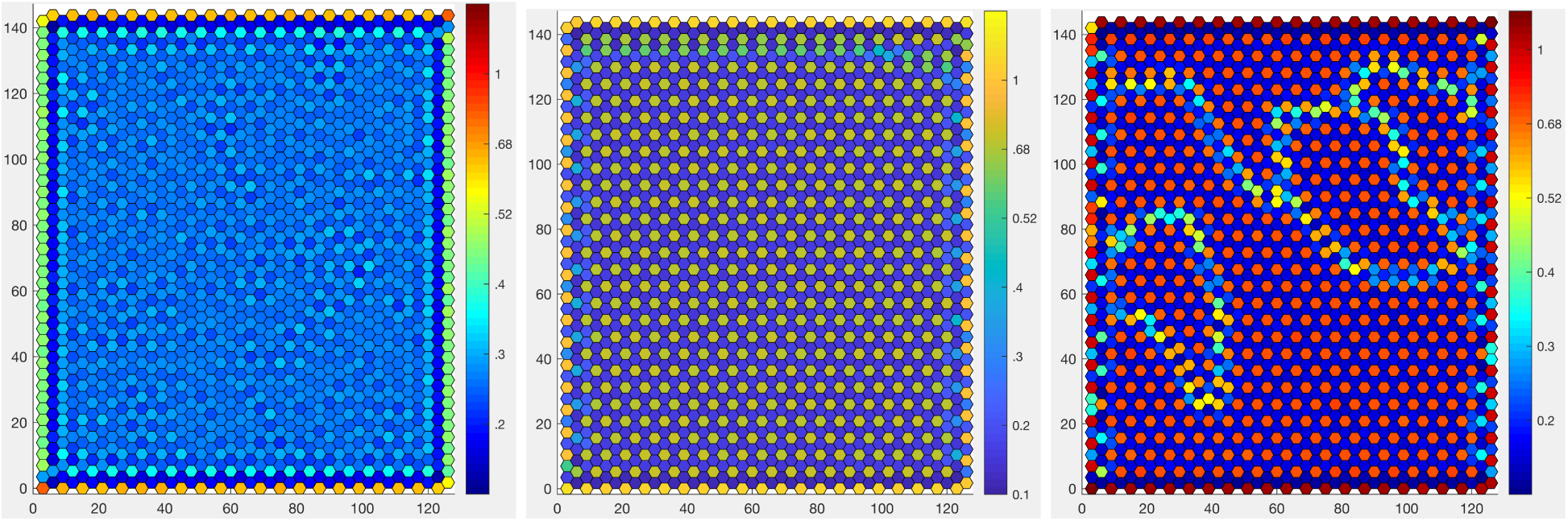
The pattern (of *D*, the patterns of *N* and *I* are similar) created by an initiation wave raising *λ_D_* from its initial value of 2 to a final value of 3.5. Left panel: The system before the arrival of the wave. Note the perturbation at the edges due to the absence of cells beyond the shown region. The bulk variation in *D* is caused by a quenched 1 percent Gaussian variation in *k*_*t*_ from cell to cell. Middle panel: The ordered pattern resulting from a slow (*v* = 0.015 cells/s) initiation wave arriving from the bottom of the system. Right panel: The multi-domain structure resulting from a fast (*v* = 0.075 cells/s) initiation wave arriving from the bottom of the system. Notice the different colormap for *D* than in Fig. 5, to better highlight the defects.

In conclusion, we have shown how the presence of a pitchfork bifurcation value 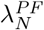 organizes the high-Delta centered hexagon pattern as well as the high-Notch centered antihexagon pattern and guarantees that the former is the generic stable structure. Futhermore, we have seen that creating a perfect pattern is a significant challenge in the vast majority of parameter space where the transition from the uniform state to the patterned state is second order. Lastly, we demonstrated that coupling a parameter to an initiation wave could provide a way to meet this challenge.

ET and DAK acknowledge the support of the United States-Israel Binational Science Foundation, Grant no. 2015619. HL acknowledges the support of the NSF grant no. PHY-1605817. ET acknowledges useful conversations with David Sprinzak.

